# Pancreatic loss of Mig6 alters murine endocrine cell fate and protects functional beta cell mass in an STZ-induced model of diabetes

**DOI:** 10.1101/2023.04.07.536046

**Authors:** Brandon M. Bauer, Jose M. Irimia, Elizabeth Bloom-Saldana, Jae-Wook Jeong, Patrick T. Fueger

**Affiliations:** Department of Molecular and Cellular Endocrinology, Arthur Riggs Diabetes and Metabolism Research Institute, City of Hope, Duarte, CA 91010, USA; Comprehensive Metabolic Phenotyping Core, Beckman Research Institute of City of Hope, Duarte, CA 91010, USA; Department of Obstetrics, Gynecology and Women’s Health, University of Missouri School of Medicine, Columbia, MO 65211

**Keywords:** errfi1, hyperglycemic clamp, dedifferentiation, proliferation, replication, glucose homeostasis

## Abstract

**Background:** Maintaining functional beta cell mass (BCM) to meet glycemic demands is essential to preventing or reversing the progression of diabetes. Yet the mechanisms that establish and regulate endocrine cell fate are incompletely understood. We sought to determine the impact of deletion of mitogen-inducible gene 6 (*Mig6*), a negative feedback inhibitor of epidermal growth factor receptor (EGFR) signaling, on mouse endocrine cell fate. The extent to which loss of Mig6 might protect against loss of functional BCM in a multiple very low dose (MVLD) STZ-induced model of diabetes was also determined.

**Methods:** Ten-week-old male mice with whole pancreas (Pdx1:Cre, PKO) and beta cell-specific (Ins1:Cre, BKO) knockout of Mig6 were used alongside control (CON) littermates. Mice were given MVLD STZ (35 mg/kg for five days) to damage beta cells and induce hyperglycemia. In vivo fasting blood glucose and glucose tolerance were used to assess beta cell function. Histological analyses of isolated pancreata were utilized to assess islet morphology and beta cell mass. We also identified histological markers of beta cell replication, dedifferentiation, and death. Isolated islets were used to reveal mRNA and protein markers of beta cell fate and function.

**Results:** PKO mice had significantly increased alpha cell mass with no detectable changes to beta or delta cells. The increase in alpha cells alone did not impact glucose tolerance, BCM, or beta cell function. Following STZ treatment, PKO mice had 18±8% higher BCM than CON littermates and improved glucose tolerance. Interestingly, beta cell-specific loss of Mig6 was insufficient for protection, and BKO mice had no discernable differences compared to CON mice. The increase in BCM in PKO mice was the result of decreased beta cell loss and increased beta cell replication. Finally, STZ-treated PKO mice had more Ins+/Gcg+ bi-hormonal cells compared to controls suggesting alpha to beta cell trans differentiation.

**Conclusions:** Mig6 exerted differential effects on alpha and beta cell fate. Pancreatic loss of Mig6 reduced beta cell loss and promoted beta cell growth following STZ. Thus, suppression of Mig6 may provide relief of diabetes.

## INTRODUCTION

Type 1 diabetes (T1D) is characterized by the severe and irreversible loss of functional beta cell mass (BCM), which results from immune-mediated depletion of beta cells and coincident loss of beta cell function due to cell stress and exhaustion [1–3]. The loss of functional BCM in T1D leads to chronic hyperglycemia and is associated with numerous comorbidities as well as shortened lifespan [4, 5]. Following clinical diagnosis of T1D, some individuals experience a so-called “honeymoon” phase, during which they experience less need for exogenous insulin. One explanation for this period is that correcting glucotoxicity with exogenous insulin allows remaining beta cells to recover and regain some functionality. However, this state is transient and is followed by near complete loss of functional BCM [6–8]. Prolonging the honeymoon period by preserving beta cell function, or even reversing beta cell loss through regeneration of beta cells, are intriguing potential treatments for T1D. Yet, gaps remain in the understanding of how beta cell mass and function are regulated during the honeymoon phase.

Several mediators of beta cell maturation, homeostasis, and growth are known. One beta cell mitogen is epidermal growth factor (EGF). Acting via its cognate receptor, EGFR, the mitogen EGF was found to aid in establishing BCM in rodents [9], and upregulation of EGF/EGFR signaling was associated with beta cell expansion [10–14]. Conversely, inhibition of EGFR has been documented to potentiate beta cell stress and death [15] Thus, in principle activating and sustaining EGFR signaling could be beneficial for fortifying functional BCM. However, the use of EGF to expand BCM has failed to advance through the therapeutic pipeline, likely due to classical feedback inhibition.

The adaptor protein, Mitogen-inducible gene 6 (Mig6; Errfi1), is one of the endogenous feedback inhibitors of EGFR [16–18]. Mig6 is indispensable for proper development and cellular homeostasis in cartilage, skin, mammary, and lung tissues [19–22]. Additionally, diabetes-associated stress such as inflammatory cytokines and glucolipotoxicity increased Mig6 gene expression [15]. Further, Mig6 has been reported to be significantly upregulated in diabetes [23]. Although Mig6 haploinsufficiency was previously reported to be protective of BCM in rodents treated with streptozotocin (STZ) [24], the specific role of Mig6 in the pancreas and the mechanisms by which it shapes BCM and beta cell function are not fully understood. We hypothesized that Mig6 is a mediator of endocrine cell homeostasis and that pancreatic loss of Mig6 would preserve BCM in a rodent model of diabetes.

Using mice in which Mig6 was selectively deleted in the whole pancreas (PKO) or beta cells (BKO), we identified that pancreatic loss of Mig6 impacted alpha and beta cell fates. Pancreatic loss of Mig6 protected mice against a multiple very low dose (MVLD) STZ paradigm of diabetes by limiting beta cell loss and promoting beta cell expansion.

## METHODS

### Animal Studies

All experiments were performed using protocols approved by the City of Hope Institutional Animal Care and Use Committee (AICUC protocol #16047) in accordance with the Guide for the Care and Use of Laboratory Animals, eighth edition [25]. Female C57Bl/6J mice harboring floxed Mig6 alleles (i.e., control Mig6^fl/fl^ mice) [26] were crossed with Mig6^fl/fl^ male mice with transgenes of either *Ins1*:Cre [27] or *Pdx*1:Cre [28]. All animals were kept under normal conditions with *ad libitum* access to food and water on standard twelve-hour light/dark cycles.

Beta cell ablation was induced with MVLD STZ (Sigma-Aldrich S0130-1G). Ten-week-old animals were administered 35 mg/kg STZ dissolved fresh in saline by intraperitoneal injection for five consecutive days as described previously [24].

For quantification of beta cell replication, starting on the third day following a complete course of STZ (35 mg/kg for 5 days), mice were treated with 100 µg/g 5-ethynyl-2’-deoxyuridine (EdU, Cayman Chemic Company CAS: 61135-33-9) by intraperitoneal injection for five consecutive days. Twenty-four hours following the final dose of EdU, mice were euthanized and pancreata were fixed, embedded in optimal cutting temperature (OCT) compound, and frozen for histology.

Intraperitoneal glucose (GTT) and insulin tolerance tests (ITT) were performed on live, conscious animals following a five-hour fast. Fasting and glucose or insulin challenged blood glucose was measured with an AlphaTRAK 2 glucometer from blood obtained from a tail nick. For glucose tolerance tests, fasted mice were administered 1.5 g/kg glucose by intraperitoneal injection. For insulin tolerance tests, fasted mice were administered 0.5 IU/kg insulin by intraperitoneal injection. Glucose readings were taken at 0, 15, 30, 60, 90, and 120 minutes after administration of glucose or insulin

### Histology

Pancreata for paraffin embedding and frozen sectioning were harvested from animals that were euthanized by CO_2_ asphyxiation. Tissues were fixed for 16 hours in 10% formalin. Following formalin fixation, pancreata were washed in 70% ethanol. With assistance of the Solid Tumor Pathology Core at City of Hope, formalin fixed paraffin embedded pancreata were sectioned into 6 µm slides. Paraffin removal was achieved by washing slides in xylene. Slides were then rehydrated in an alcohol gradient from 100 – 70%. For antigen retrieval, samples were immersed in boiling citric acid buffer (Vector Labs, H3300) for 15-30 minutes and allowed to cool to room temperature. Tissues were blocked with either serum free protein blocking reagent (Dako Agilent X0909) or 5% BSA. Antibodies were diluted in either background reducing antibody diluent (Dako Agilent S3022) or 2% BSA. Antibodies used were guinea pig anti insulin (Abcam ab195956, 1:500), mouse anti glucagon (Abcam ab10988, 1:500), rabbit anti somatostatin (Thermo Fisher Sci. PA5-82678, 1:500) rabbit anti ALDH1A3 (Novus Biologicals NBP2-15339, 1:500), rabbit anti MAFA (Bethyl Laboratories BLR067G, 1:200), and Invitrogen Alexa Fluor conjugated secondary antibodies produced in goat. For immunohistochemistry, secondary biotinylated goat anti guinea pig (Vector BA7000, 1:500) or biotinylated goat anti mouse (1:500) antibodies were used, followed by Vectastain Elite ABC HRP detection kit (Vector PK-6100) and detection with DAB (Agilent Technologies K346811-2). Cell death and DNA fragmentation were detected with a TUNEL In Situ Cell Death Detection Kit (Roche, Cat. 12156792910). Frozen pancreata for histology were removed and fixed as described above. Fixed pancreata were then incubated overnight in 30% sucrose and then embedded in OCT. Slides were sectioned at 6 µM on a Leica CM3050 S cryostat (Leica Biosystems; Deer Park, IL) and stored at −80°C until use. EdU detection was carried out using the Click-iT EdU 488 Imaging Kit (Invitrogen, C10337) according to the manufacturer’s instructions.

### Microscopy and Image Analysis

For immunohistochemistry analysis, 2-3 slides per animal with >50 µm of separation between slides were analyzed. The relative beta cell area was determined as the average of the beta cell areas. Images were obtained using Keyence BZ-II all-in-one microscope (Keyence Corporation of America; Itasca, IL) and analyzed with Keyence software.

Immunofluorescent images were obtained using a Zeiss Axio Observer 7 with ApoTome.2 for widefield deconvolution. Immunofluorescent image analysis was performed in QuPath (version 0.3.2) with assistance from the Light Microscopy Core at City of Hope. A minimum of 12 islets from at least two different slides were used per experiment unless otherwise stated. Islets had to contain more than 10 visible endocrine cells to be included in the analysis. All analysis was performed in a blinded fashion.

### Islet Isolation and Culturing

Islets were isolated from 10-week-old male mice using established methods [29]. Mice were euthanized by CO_2_ asphyxiation. Pancreata were inflated with 2.5 ml CIZyme RI (Vitacyte, Cat. #005-1030) prepared according to manufactures instructions. Isolated islets were picked from cellular debris and cultured overnight at 37°C, with 5% CO_2_ in RPMI media supplemented with 10% FBS, 10mM HEPES, and penicillin-streptomycin.

For protein quantification from isolated islets, a minimum of 50 islets were lysed in radioimmunoprecipitation assay (RIPA) buffer. Protein (750 ng) was then loaded into individual wells of a Protein Simple Wes cassette, and samples were run according to manufacturer instructions. Antibodies used included Actin (MP Biomedicals 691001, 1:100) and ALDH1A3 (Novus Biologicals NBP2-15339, 1:100).

### qPCR and mRNA Sequencing

A minimum of 75 islets harvested from a single mouse were lysed in RLT buffer with 0.1% 2-Mercaptoethanol. RNA was isolated using RNeasy Micro (Qiagen). RNA concentration and purity were confirmed using Nanodrop. From isolated RNA, cDNA libraries were constructed for qPCR analysis using Applied Biosystems High-Capacity cDNA Reverse Transcription Kit (Fisher Scientific #4375575). Quantification with QuantStudios 7 Flex Real-Time PCR system and Taqman probes (Thermo). RNA sequencing libraries were prepared with Kapa RNA mRNA HyperPrep kit (Kapa Biosystems, Cat KR1352) according to the manufacturer’s protocol. The final libraries were validated with the Agilent Bioanalyzer DNA High Sensitivity Kit and quantified with Qubit. Sequencing was performed on Illumina NovaSeq 6000 by using S4 kit v1.5 with the paired end mode of 2×101cycle. Real-time analysis (RTA) v3.4.4 software was used for base calling.

### Statistical Analysis

Data are presented as means ± SEM. Protein and mRNA data were normalized to control conditions and presented as relative expression. Adjusted area-under-the-curve (AUC) was calculated using the trapezoid method. Student’s *t*-test (unpaired, two-tailed unless otherwise stated) or ANOVA (with Bonferroni post-hoc tests) were performed using GraphPad Prism software (La Jolla, CA, USA) to detect statistical differences. *p < 0.05* was considered statistically significant.

## RESULTS

### Loss of Mig6 promotes alpha cell hypertrophy

We previously demonstrated that reduced levels of Mig6 mitigated beta cell damage and promoted beta cell proliferation in response to stress [30–32]. To expand on these results and determine tissue specificity of the previous results, the effect of Mig6 loss on endocrine cell fate and homeostasis in adult mice was determined. To this end, Mig6 knock-out mouse lines in which Mig6 was constitutively knocked out of either the whole pancreas (pdx1:Cre, PKO) [28] or from beta cells alone (ins1:cre, BKO) were employed [27]. We confirmed loss of Mig6 gene expression by qRT-PCR from isolated islets from PKO, BKO, or littermate controls (CON; **Figure 1A**). At ten weeks of age, neither the PKO mice nor the BKO mice had significant differences in body or pancreas weight compared to CON mice (**Figure 1B, C, Supplementary Figure 1**).

**Figure 1:**
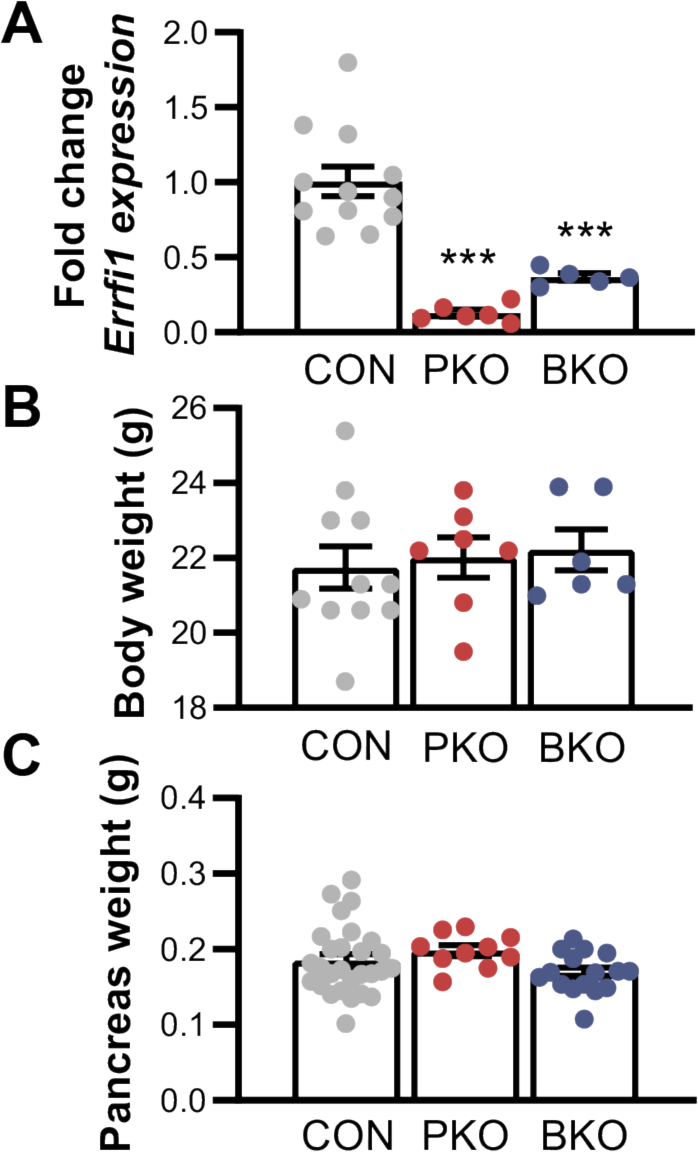
Whole pancreas or beta cell-specific deletion of mitogen-inducible gene 6 does not change pancreas or body weight. **A**. *Errfi1* (Mig6) gene expression in islets isolated from 10-week-old mice was quantified by RT-PCR. Expression was significantly decreased in PKO and BKO animals (n >5 mice). **B.** Animal body and pancreas (**C**) weight were determined (n >6 animals per strain). ANOVA; p-values: * <0.05, ** <0.01, *** <0.001; data reported as means ± SEM.

To determine the impact of Mig6 loss on endocrine cell fate, we labeled pancreas sections from ten-week-old male mice with the markers of pancreatic endocrine cells: glucagon positive alpha cells, insulin positive beta cells, and somatostatin positive delta cells (**Figure 2A**). We detected significant differences in the endocrine cell ratios between PKO and CON animals (**Figure 2B**). As a percent of the total islet, PKO mice had a significant decrease in the relative insulin positive area and a significant increase in the relative glucagon positive area compared to CON mice. Interestingly, differences in the distribution of endocrine cells between the CON and BKO animals were not observed.

**Figure 2:**
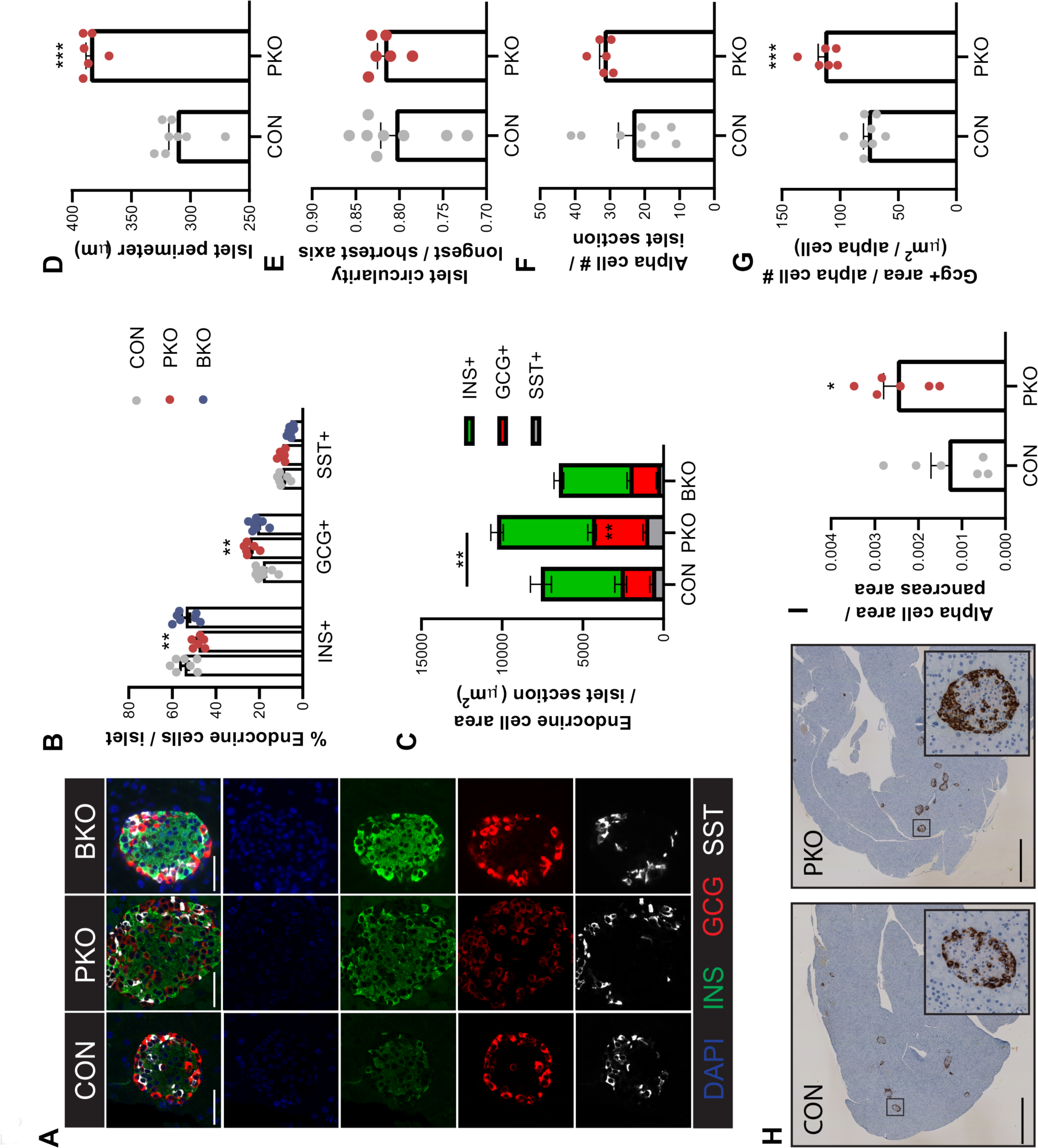
Pancreatic loss of Mig6 alters endocrine cell fate without impacting functional beta cell mass. **A.**Representative immunofluorescence images of pancreas tissue sections from 10-week-old mice show the distribution of beta cell (INS+), alpha cell (GCG+), and delta cell (SST+) positive area. **B.** Data was quantified as a percent of total islet area. **C.** Raw area (µm^2^) of each endocrine cell type, which collectively yielded the total islet size. **D.** Islet perimeter was independently quantified. **E.** Islet circularity was calculated as the ratio of the longest to shortest axis. **F.** Alpha cell number was manually counted. **G.** Alpha cell size, calculated as the total alpha positive area / # alpha cells per islet cross-section. **H-I.** Glucagon positive area / pancreas area was determined by IHC labeling of whole pancreata. n = 6-8 mice; ANOVA or 2-tailed student t-test; p values: * <0.05, ** <0.01, *** <0.001; data reported as means ± SEM.

Given the differences in the relative percentages of endocrine cells, we compared absolute endocrine cell area of each endocrine cell type. A 70.3±13.5% increase in alpha cell area in PKO mice compared to CON was identified. There was no significant difference in either beta or delta cell area between the knockout and CON mice (**Figure 2C**). In contrast to PKO mice, alpha cell area of BKO animals was indistinguishable from CON. These results suggested that an increase in the absolute area of alpha cells accounted for the relative difference in beta cells described in Figure 2B. The increase in alpha cell area also increased the overall islet area in the PKO mice compared to CON mice. To validate the increase in islet size, we quantified islet perimeter. In line with an expansion of endocrine cells, a significant increase in the islet perimeter was noted in organs from PKO mice compared to CON mice (**Figure 2D**). Whereas the size of islets was increased, there were no changes in islet shape, as measured by circularity (**Figure 2E**).

To determine the source of increased alpha cell area, we counted the number alpha cells per islet section. The alpha cells tended to be more numerous in PKO islets, but this difference was not quite significant (p=0.0529; **Figure 2F**). There was, however, a significant increase in alpha cell size (glucagon positive area/alpha cell number), suggesting that pancreatic loss of Mig6 resulted in alpha cell hypertrophy (**Figure 2G**). The alpha cell hypertrophy was confirmed by quantification of the glucagon positive area in pancreata from the respective mice (**Figure 2H, I**).

### Pancreatic loss of Mig6 is protective against MVLD-STZ induced diabetes but loss from beta cells alone is insufficient for protection

Further analysis of pancreata under normal conditions confirmed similar beta cell area regardless of Mig6 status (**Figure 3A-B**). However, it was not clear if loss of Mig6 impacted islet function. Fasting blood glucose was similar between in BKO, PKO, and CON mice. As well, all groups exhibited the same degree of glucose tolerance under normal conditions (**Figure 3C-D**). We previously described how Mig6 haploinsufficiency protected against STZ-induced diabetes [24]. To identify which cellular compartments may contribute to these beneficial effects, young PKO, BKO, and CON mice were treated with 35 mg/kg STZ for five days to induce beta cell dysfunction and death (**Figure 4A**). Twenty days later, whereas both BKO and CON mice presented with significantly increased fasting blood glucose and impaired glucose tolerance, PKO mice had normal fasting blood glucose and were partially protected from impaired glucose tolerance (**Figure 4B-C**). Consistent with the improved glucose tolerance, PKO mice had

**Figure 3:**
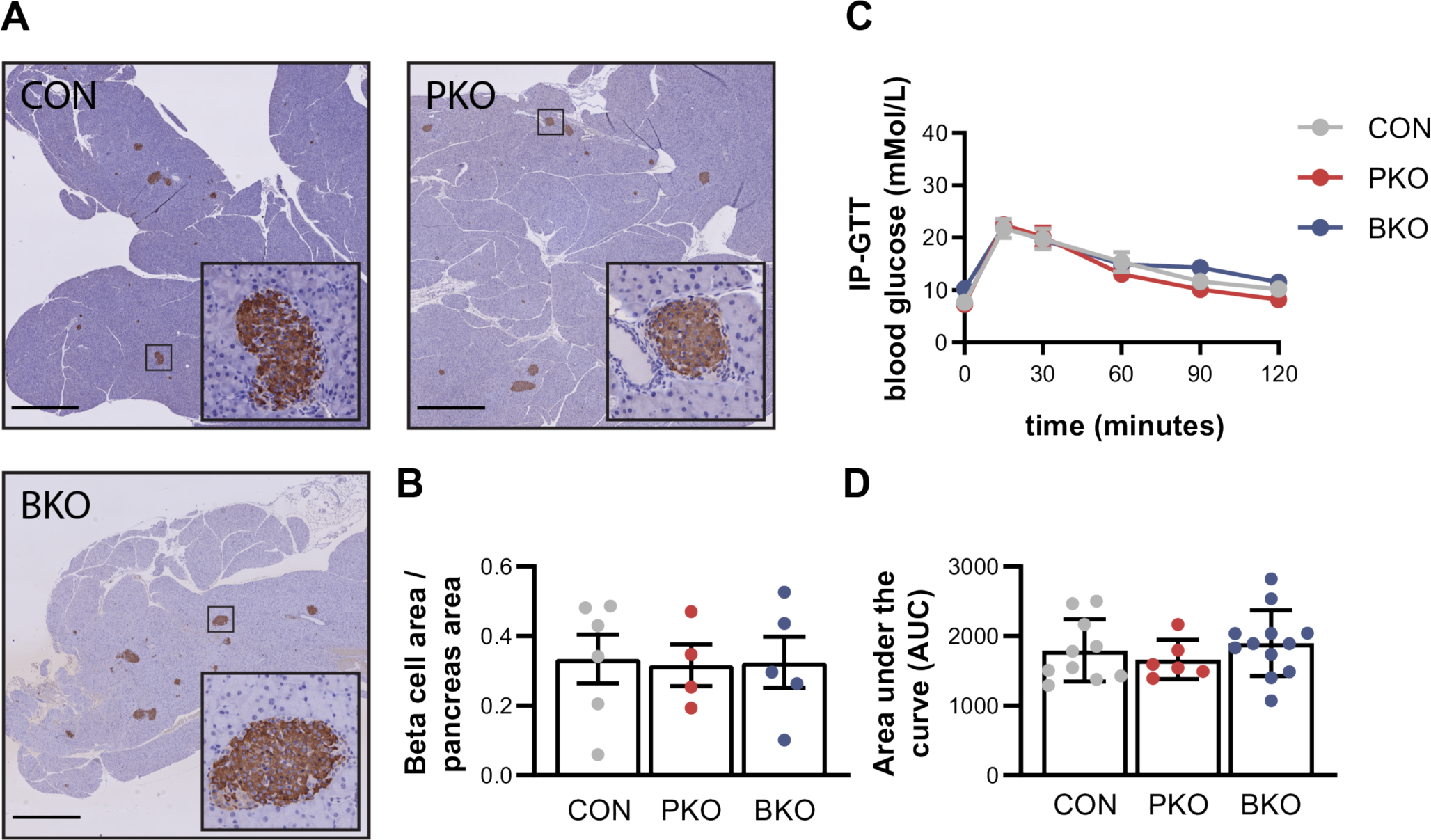
Pancreatic loss of Mig6 does not alter beta cell function or glucose tolerance in non-diabetic mice. **A-B.** Representative images and quantification of pancreatic Ins+ beta cell area as demarcated by IHC labeling of whole pancreata from CON, PKO, and BKO animals. **C.** 10-week-old male mice were challenged with a 1.5 g/kg glucose bolus i.p. and blood glucose levels determined. **D.** Calculated area-under-the-curve (AUC). N = 7-12; ANOVA, p values: * <0.05, ** <0.01, *** <0.001; data reported as means ± SEM.

**Figure 4:**
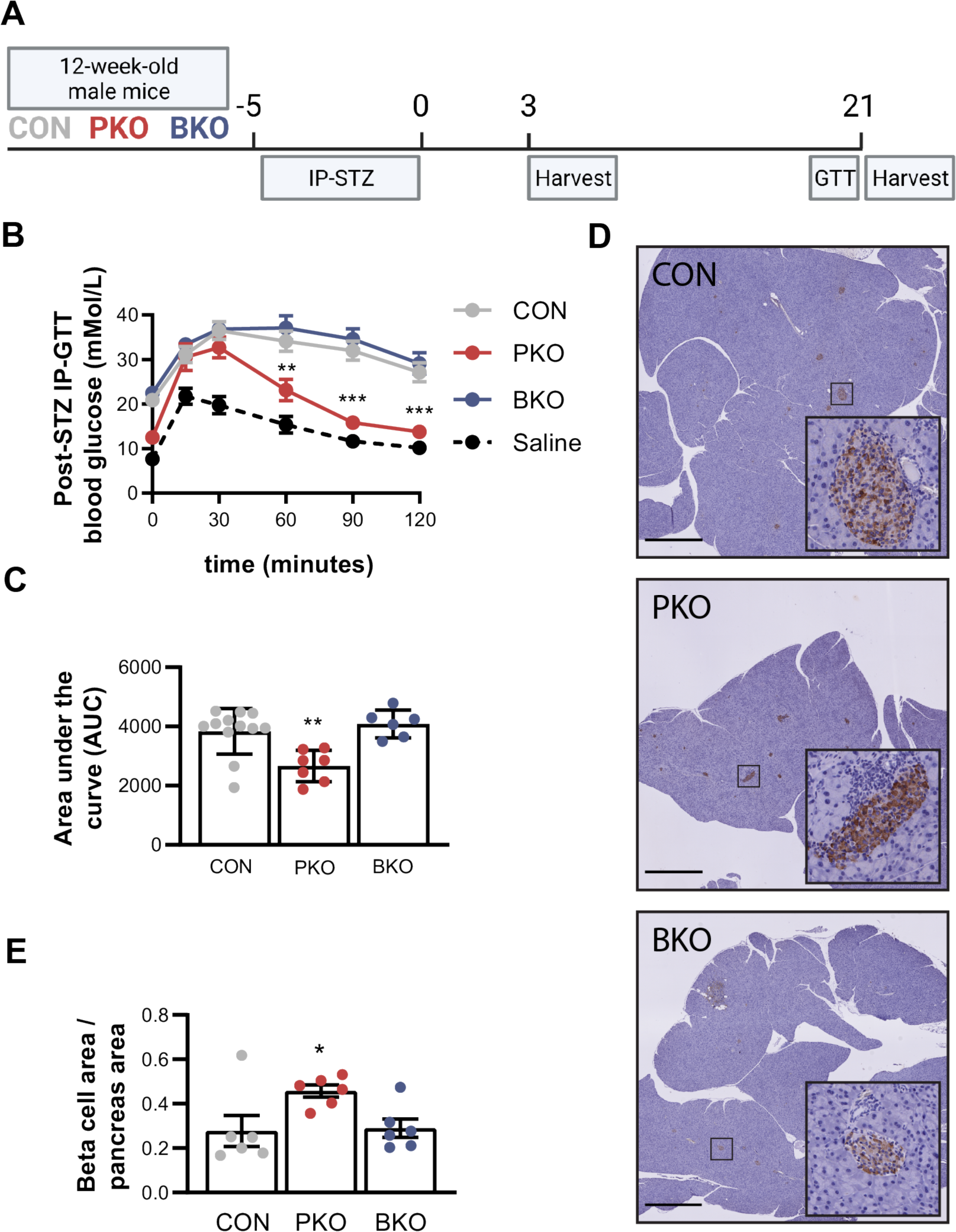
Beta cell-specific loss of Mig6 does not protect against STZ-induced diabetes. **A**. Pancreas and beta cell-specific null mice were administered STZ (35 mg/kg in saline i.p.) for 5 consecutive days. A glucose tolerance test was performed on the third and twentieth day following STZ. Isolated islets or whole pancreata were harvested on days 3 and 21 after STZ. **B.** Blood glucose levels were measured in mice after a glucose bolus of 1.5 g/kg i.p. **C.** AUC was calculated. **D-E.** Representative images and quantification of pancreatic Ins+ beta cell area determined by IHC labeling of whole pancreata of CON, PKO, and BKO mice. n = 6-12; ANOVA, p values: * <0.05, ** <0.01, *** <0.00; data reported as means ± SEM.

64.9±9.6% greater BCM than CON animals (**Figure 4D-E**).

### Loss of Mig6 mitigates beta cell stress without impacting cell death

Beta cell specific cytotoxicity induced by MVLD-STZ causes mice to experience islet cell inflammation, endoplasmic reticulum stress, beta cell dysfunction, and cell death [33–35]. Overexpression of Mig6 was previously found to exacerbate ER stress and apoptosis in rat insulinoma cell lines [30]. From isolated islets of STZ treated mice, we determined that loss of Mig6 decreased stress-induced *Ddit3* expression. In contrast, Mig6 loss upregulated genes associated with p53-mediated apoptotic pathways (**Figure 5A**). Nonetheless, after STZ, the degree of TUNEL staining, marking apoptotic cell death, did not vary appreciably between islets from CON or PKO mice (**Figure 5B-C**).

**Figure 5:**
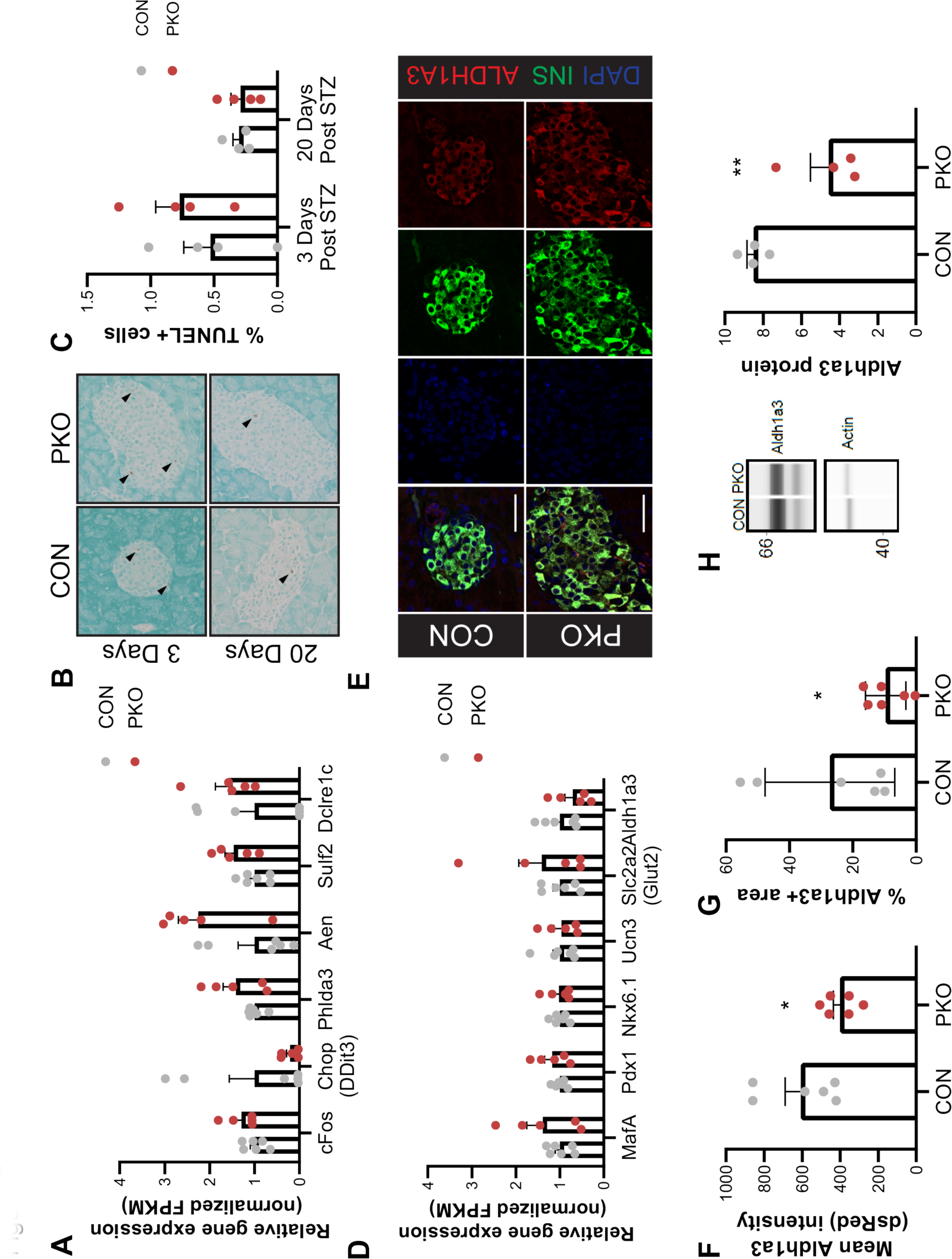
Loss of Mig6 reduces STZ-induced beta cell loss and beta cell dedifferentiation. **A.** Beta cell stress, apoptosis, and death-associated gene mRNA levels were determined in isolated islets from 10-week-old CON, PKO, and BKO mice on the third day following STZ. Data shown as relative FPKM values normalized to CON. **B-C**. Cell death was assessed by labeling TUNEL+ cells before and on days 3 and 21 following STZ. **D.** Endocrine cell identity and maturity associated gene mRNA levels were assessed in isolated islets by RNA seq. Data shown as relative FPKM values normalized to CON. **E.** Representative images of Aldh1a3+ cells dedifferentiation cells. **F.** Quantification of protein expression by immunofluorescent intensity. **G.** The area above a threshold in whole pancreata from CON and PKO mice. **H-I.** Protein from isolated islets from STZ-treated mice was quantified by Protein Simple Wes capillary-based quantification. Shown are representative false color images and quantification of protein bands relative to actin controls. n = 4-6; 2-tailed t-test, p values: * <0.05 ** <0.01 *** <0.001; data reported as means ± SEM.

### PKO mice have decreased beta cell dedifferentiation following STZ treatment

High dose STZ treatment induces near complete ablation of beta cells [33], whereas MVLD-STZ, while less toxic to cells, still promoted hyperglycemia in mice likely through effects on beta cell fate and or function [34]. In keeping with this, loss of Mig6 tended to maintain or increase expression of beta cell maturity genes such as *MafA, Pdx1,* and *Slc2a2* while decreasing *Aldh1a3* a beta cell dedifferentiation gene (**Figure 5D**). Mice exposed to MVLD-STZ are reported to exhibit increased *Aldh1a3* [34, 36]. Indeed, mice exposed to MVLD-STZ had increased *Aldh1a3* (data not shown) although the degree of increase was less pronounced in PKO mice (**Figure 5E-G**). Further, islets isolated from PKO mice had lower levels of ALDH1A3 protein compared to islets from CON mice (**Figure 5H**). Other markers of beta cell maturity and senescence including PDX1, MAFA, and P21 did not vary by levels of Mig6 (**Supplementary Figure 2**). ALDH1A3 is an early marker of dedifferentiation [36, 37]. Therefore, Mig6 may enforce beta cell maturation and exit from the cell cycle, all to preserve beta cell functionality.

### Loss of Mig6 promotes beta cell replication following STZ

BCM was retained in PKO mice and even increased after STZ treatment (**Figure 3A-B, 4D-E**). The increase in BCM may occur secondary to transdifferentiation of mature endocrine cells [38], neogenesis from progenitor cells [39], and/or beta cell proliferation [40–42]. mRNA levels of several cell cycle-and proliferation-related genes (*Cdk1*, *Wee1*, *Ccnb1*, and *Rprd1b*) were increased in islets from PKO mice (**Figure 6A**). Beta cell replication is minimal in adult mice and decreases further with age [43, 44]. To test if Mig6 deletion altered beta cell proliferation, mice were treated for five days with 5-ethynyl-2’-deoxyuridine (EdU) beginning on the third day following the MVLD-STZ treatment and euthanized on the eighth day. Pancreata were then evaluated for EdU incorporation by immunofluorescence (**Figure 6B**). PKO mice had significantly higher numbers of EdU-positive cells compared to CON mice. Of interest, BKO mice had similar rates of EdU-positive endocrine cells compared to CON mice (**Figure 6C-E**) suggesting that beta cell-specific loss of Mig6 is insufficient to promote beta cell proliferation after STZ injury.

**Figure 6:**
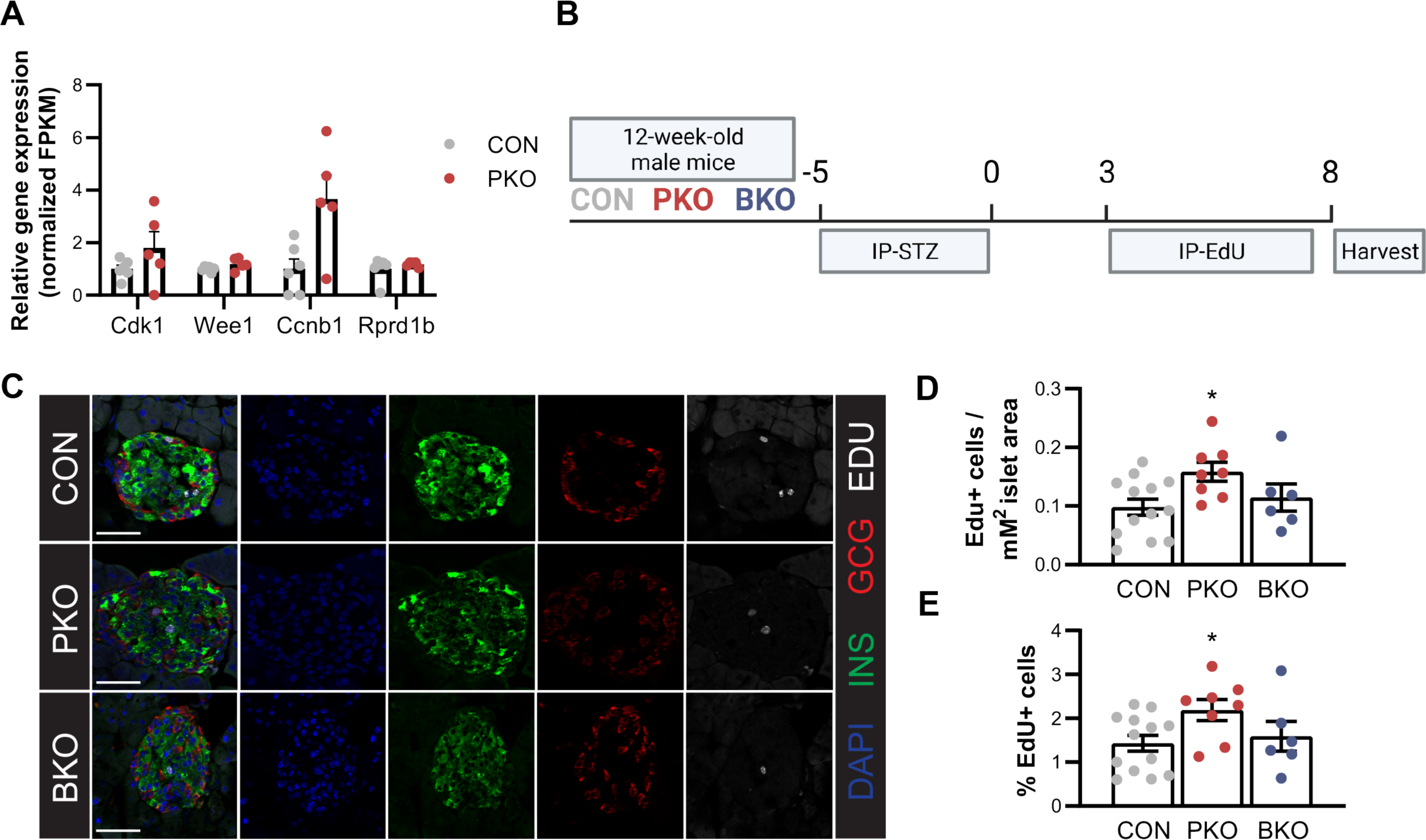
Mig6 mediates of beta cell expansion following STZ-induced beta cell injury. **A.** Genes associated with cell cycle progression and proliferation were quantified by RNA Seq. Data shown as relative FPKM values normalized to CON. **B**. To quantify beta cell replication, mice were treated for 5 days with STZ and on the third day following STZ were administered 100 µg/g Edu. **C** Edu+ beta cells were identified by immunofluorescence. **D-E** Data were quantified as the total number of cells per islet as well as the percent of Edu+ beta cells. n = 4-13 2-way ANOVA, p values: * <0.05 ** <0.01 *** <0.001, data reported as means ± SEM.

### Loss of Mig6 may promote alpha to beta cell transdifferentiation

PKO mice had more EdU-positive islet cells after STZ. However, the proliferative rate was still low (∼2% of cells) (**Figure 6**). The low, albeit increased, proliferation rate raised the notion that the enhanced BCM in PKO mice could be driven through other mechanisms such as transdifferentiation. Exploring this possibility, 21 days after STZ treatment, islet alpha cell area was now normalized in PKO mice compared to CON mice (**Figure 7A-B**), in contrast, the expanded alpha cell area in PKO compared to CON mice before STZ treatment previously discussed (**Figure 2**). To further examine these results, pancreas sections were labelled with markers of endocrine cells: insulin (beta cells), glucagon (alpha cells), and somatostatin (delta cells) (**Figure 2C**). Supportive of the IHC data, we detected no increase in alpha cell area between the CON and KO mice following treatment with STZ (**Figure 7D**). Islet cells expressing both insulin and glucagon, so-called Ins+/Gcg+ bihormonal cells, are believed to be indicative of cellular transdifferentiation [45, 46]. Prior to STZ, bihormonal cells were not detected in islets from mice (data not shown and **Figure 2**). However, with STZ exposure, islets from CON and PKO mice had bihormonal endocrine cells. The bihormonal cells persisted at 3 weeks after STZ in PKO but not CON mice (**Figure 7E-F**).

**Figure 7:**
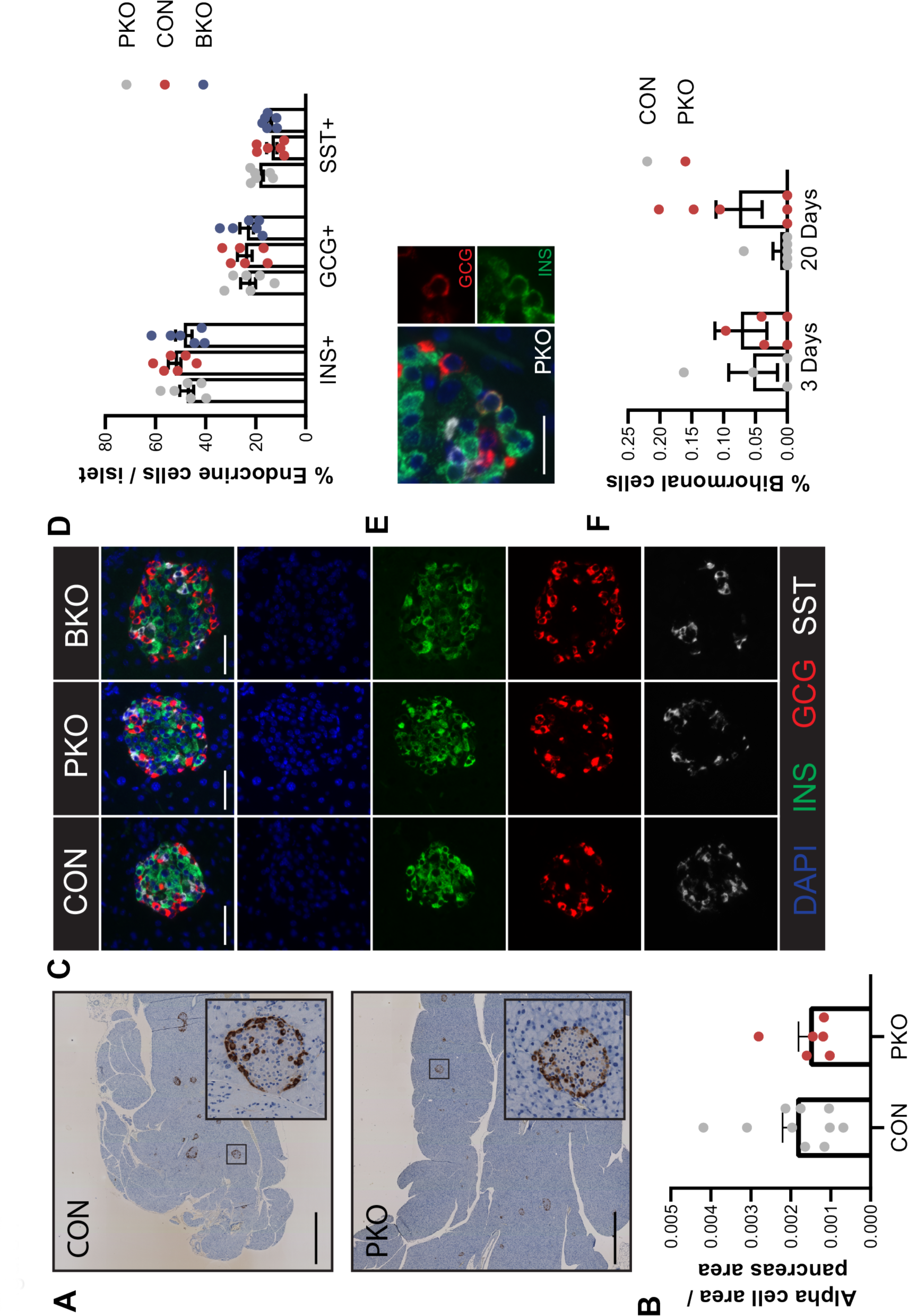
Loss of Mig6 may promote alpha to beta cell transdifferentiation. **A**. Representative images of alpha cell area in animals 21 days after treatment with STZ as determined by IHC. **B.** Quantification of alpha cell area. **C.** Representative images of insulin (beta cell), glucagon (alpha cell), and somatostatin (delta cell) positive area assessed by immunofluorescence. **D.** Quantification of endocrine cell percentages. **E-F**. Representative image of an insulin positive/glucagon positive bihormonal cell and quantification of bihormonal cells observed in PKO and CON mice. n = 4-10 2-way ANOVA or 2-tailed t-test; p values: * <0.05 ** <0.01 *** <0.001; data reported as means ± SEM.

## DISCUSSION

The regulation of endocrine cell fate and the mechanisms that protect BCM in diabetes remain poorly understood. EGFR signaling stimulates beta cell formation and expansion [9, 14, 47], and thus has been explored as a therapeutic target for the treatment of diabetes [11, 12, 48, 49]. However, successes in pre-clinical models have failed to translate to human diabetes. We posit that feedback inhibition of EGFR by Mig6 limits the therapeutic efficacy of this pathway. Herein, we identified that Mig6 ablation altered alpha and beta cell fates under normal conditions and in response to beta cell stress and destruction imposed by STZ. Additionally, in mice rendered hyperglycemic by STZ, pancreatic loss of Mig6 preserved functional BCM and mitigated hyperglycemia. In PKO mice, BCM maintenance was likely attributed to the combined effects of decreased beta cell loss and increased beta cell proliferation; endocrine cell transdifferentiation might have also contributed to the preservation of BCM following STZ treatment. Given the broad effects of EGFR signaling, it is not surprising that Mig6 deletion could impact multiple processes involved in the preservation and expansion of beta cell mass.

Mig6 deletion was also found to more broadly influence endocrine cell size and number, especially in alpha cells. Islets from PKO mice were larger with more alpha cell mass. This latter finding was mostly due to cell hypertrophy, although it is possible an effect on alpha cell hyperplasia also exists. Other endocrine cell types examined, however, seemed to be unaffected by pancreatic Mig6 deletion. This expanded alpha cell pool appeared to serve as a reservoir of cells that could be exploited to transdifferentiate into beta cells when functional beta cell mass decreased. Indeed, many reports have previously demonstrated alpha-to-beta cell transdifferentiation [50–52]. Here, whereas we did not perform lineage tracing studies, the gold standard for detecting alpha-to-beta transdifferentiation [53, 54], we did observe that the expanded alpha cell area was depleted by STZ, which coincided with a preservation and even increase in BCM after STZ treatment.

In contrast to pancreatic Mig6 deletion, beta cell-specific Mig6-null islets mimicked control islets regarding the distribution of endocrine cells. These data suggest that the early deletion of Mig6 in PDX-1 positive, endocrine precursor cells during development led to the alterations in cell fate. By deleting Mig6 in insulin positive cells later in development, the opportunities for cell fate switching or expansion are more limited. Importantly, PKO mice showed indications of beta cell growth following STZ. Interestingly, with time, the post-STZ increase in alpha cell mass dissipated. However, whereas an increase in insulin positive, glucagon positive bihormonal cells, which is thought to be indicative of transdifferentiating cells, was detected, this occurrence was somewhat uncommon. A likely explanation for the limited appearance of bihormal cells is that it represents a short-lived, transient state. Together these observations support, but do not completely demonstrate, alpha-to-beta cell transdifferentiation. Future studies using lineage tracing models could provide definitive evidence.

Mig6 participates in the development and homeostasis of several tissues [19–22]. Complete loss of Mig6 is associated with increased embryonic lethality and significantly decreased life span in mice [19, 26]. Mig6 governs cell migration, branching morphogenesis, proliferation, senescence, and apoptosis, although the exact roles of Mig6 are cell- and tissue-specific [55]. Prior to this report, the comprehensive actions of Mig6 in the pancreas were poorly understood. Further, the previous work on Mig6 has largely supported a cell autonomous role for Mig6. Thus, it was our expectation that deletion of Mig6 in the beta cells would phenocopy the results of the PKO model. Surprisingly, these beneficial effects of Mig6 deletion did not extend to the beta cell-specific knockout model, suggesting that some mechanism of endocrine-endocrine or exocrine-endocrine crosstalk exist to drive the beneficial effects reported in the PKO mice. Deletion of Mig6 selectively in other endocrine cells (e.g., alpha cells), exocrine, or ductal cells could provide evidence for such a crosstalk mechanism.

Despite our advancements in defining the role of Mig6 in the pancreas, this study has some caveats worth noting. First, the timing of Mig deletion likely influenced the phenotypic results reported herein. That is, deleting Mig6 beyond a critical developmental window with the use of inducible mouse models may alter how the endocrine cells respond during normal and stressed conditions, such as when challenged by STZ. Second, how readily these results translate to human islets is unknown. Once available, Mig6 inhibitors or deletion strategies could be used to examine the impact on human beta cell proliferation, dedifferentiation, and alpha-to-beta cell transdifferentiation. Third, by default, it is assumed that most of the responses observed in the PKO mice reported here are due to heightened or prolonged signaling of EGFR or other Erbb/HER family members. However, as an adapter protein, Mig6 could exert effects independent of EGFR. Indeed, we have recently reported an interaction between Mig6 and the adaptor protein NumbL that controls beta cell death during glucolipotoxicity [56]. Mig6 can also exert effects through direct interaction with other proteins, including another adaptor protein SHC1 [57], the RhoGTPase Cdc42 [58], WEE1 of the cell cycle machinery [59], among others. Finally, STZ creates a model of beta cell destruction, resulting in insulin deficiency. Whereas this model exhibits some features central to human T1D, one needs to be careful when extrapolating these results to the human disease.Nevertheless, a recent report indicated that *Errfi1* (human Mig6 gene) is among the most upregulated genes in T1D [23]. Therefore, it would be useful to carefully interrogate the signatures of EGRF and Mig6 in islet endocrine cells as well as other cell types from of individuals with T1D.

In summary, pancreatic loss of Mig6 mitigated the effects of low dose STZ on glucose homeostasis, promoted beta cell growth, and may have initiated islet endocrine cell transdifferentiation. However, similar benefits were not provided by beta cell specific Mig6 deletion. Whereas islet endocrine cell deletion of Mig6 appeared important, potential effects upon non-endocrine and vascular cells in pancreas null animals should be considered. Thus, Mig6 should be considered as a therapeutic target for preserving functional beta cell mass as part of combination therapy combining immunomodulatory agents and/or beta cell mitogens.

## Acknowledgements

This work was supported by a National Institutes of Health grant DK099311 to PTF and Innovation Award from the Wanek Family Project to Cure Type 1 Diabetes from the City of Hope to PTF. Research reported in this publication included work performed in City of Hope’s Comprehensive Metabolic Phenotyping Core and the Solid Tumor Pathology, Integrated Genomics, and Light Microscopy Cores as well as Center for Comparative Medicine, which were supported by the National Cancer Institute of the National Institutes of Health under grant number P30CA033572. The content is solely the responsibility of the authors and does not represent the official views of the National Institutes of Health. Thanks to Dr. Dustin Schones and Michael Nelson for support of RNA-seq and image analyses, respectively. Finally, thanks to Dr. Jeffrey Isenberg for help with editing the manuscript.

## Author Contributions

Author contributions: conceptualization, BMB and PTF; methodology, BMB, JI, and PTF; formal analysis, BMB, JI, EB-S, and PTF; investigation, BMB, JI, EB-S, and PTF; resources, J-WJ and PTF; data curation, BMB; writing - original draft preparation, BMB and PTF; writing - review and editing, BMB, JI, EB-S, J-WJ, and PTF; visualization, BMB; supervision, PTF; project administration, BMB and PTF.; funding acquisition, PTF. All authors have read and agreed to the published version of the manuscript.

## Funding

This research was funded by National Institutes of Health grant DK099311 to PTF and Innovation Awards from the Wanek Family Project to Cure Type 1 Diabetes to PTF. This work utilized services from the Center for Comparative Medicine and Solid Tumor Pathology, Integrated Genomics, and Light Microscopy Cores at the City of Hope, which were supported by the National Cancer Institute of the National Institutes of Health grant P30CA033572.

## Conflicts of Interest

The authors declare no conflicts of interest.

## Notes

### Competing Interest Statement

The authors have declared no competing interest.

## REFERENCES

[1] Roep BO, Thomaidou S, van Tienhoven R, Zaldumbide A (2021) Type 1 diabetes mellitus as a disease of the β-cell (do not blame the immune system?). Nature Reviews Endocrinology 17(3): 150–161. 10.1038/s41574-020-00443-4

[2] Chen C, Cohrs CM, Stertmann J, Bozsak R, Speier S (2017) Human beta cell mass and function in diabetes: Recent advances in knowledge and technologies to understand disease pathogenesis. Molecular Metabolism 6(9): 943–957. https://doi.org/10.1016/j.molmet.2017.06.019

[3] Katsarou A, Gudbjornsdottir S, Rawshani A, et al. (2017) Type 1 diabetes mellitus. Nature reviews Disease primers 3: 17016. 10.1038/nrdp.2017.16

[4] Copenhaver M, Hoffman RP (2017) Type 1 diabetes: where are we in 2017? Translational Pediatrics 6(4): 359–364. 10.21037/tp.2017.09.09

[5] de Ferranti SD, de Boer IH, Fonseca V, et al. (2014) Type 1 Diabetes Mellitus and Cardiovascular Disease: A Scientific Statement From the American Heart Association and American Diabetes Association. Diabetes Care 37(10): 2843

[6] van Megen KM, Spindler MP, Keij FM, et al. (2017) Relapsing/remitting type 1 diabetes. Diabetologia 60(11): 2252–2255. 10.1007/s00125-017-4403-3

[7] Tang R, Zhong T, Wu C, Zhou Z, Li X (2019) The Remission Phase in Type 1 Diabetes: Role of Hyperglycemia Rectification in Immune Modulation. Frontiers in Endocrinology 10

[8] Abdul-Rasoul M, Habib H, Al-Khouly M (2006) ‘The honeymoon phase’ in children with type 1 diabetes mellitus: frequency, duration, and influential factors. Pediatric Diabetes 7(2): 101–107. https://doi.org/10.1111/j.1399-543X.2006.00155.x

[9] Miettinen P, Ormio P, Hakonen E, Banerjee M, Otonkoski T (2008) EGF receptor in pancreatic β-cell mass regulation. Biochemical Society Transactions 36(3): 280–285. 10.1042/bst0360280

[10] Maachi H, Fergusson G, Ethier M, et al. (2020) HB-EGF Signaling Is Required for Glucose-Induced Pancreatic β-Cell Proliferation in Rats. Diabetes 69(3): 369. 10.2337/db19-0643

[11] Suarez-Pinzon WL, Yan Y, Power R, Brand SJ, Rabinovitch A (2005) Combination therapy with epidermal growth factor and gastrin increases beta-cell mass and reverses hyperglycemia in diabetic NOD mice. Diabetes 54(9): 2596–2601

[12] Song I, Patel O, Himpe E, Muller CJF, Bouwens L (2015) Beta Cell Mass Restoration in Alloxan-Diabetic Mice Treated with EGF and Gastrin. PLOS ONE 10(10): e0140148. 10.1371/journal.pone.0140148

[13] Hakonen E, Ustinov J, Palgi J, Miettinen PJ, Otonkoski T (2014) EGFR Signaling Promotes β-Cell Proliferation and Survivin Expression during Pregnancy. PLOS ONE 9(4): e93651. 10.1371/journal.pone.0093651

[14] Zarrouki B, Benterki I, Fontés G, et al. (2014) Epidermal growth factor receptor signaling promotes pancreatic β-cell proliferation in response to nutrient excess in rats through mTOR and FOXM1. Diabetes 63(3): 982–993. 10.2337/db13-0425

[15] Chen Y-C, Lutkewitte AJ, Basavarajappa HD, Fueger PT (2020) Glucolipotoxicity-induced Mig6 desensitizes EGFR signaling and promotes pancreatic beta cell death. bioRxiv: 2020.2011.2024.380279. 10.1101/2020.11.24.380279

[16] Xu D, Makkinje A, Kyriakis JM (2005) Gene 33 is an endogenous inhibitor of epidermal growth factor (EGF) receptor signaling and mediates dexamethasone-induced suppression of EGF function. The Journal of biological chemistry 280(4): 2924–2933. 10.1074/jbc.M408907200

[17] Anastasi S, Lamberti D, Alema S, Segatto O (2016) Regulation of the ErbB network by the MIG6 feedback loop in physiology, tumor suppression and responses to oncogene-targeted therapeutics. Seminars in cell & developmental biology 50: 115–124. 10.1016/j.semcdb.2015.10.001

[18] Zhang X, Pickin KA, Bose R, Jura N, Cole PA, Kuriyan J (2007) Inhibition of the EGF receptor by binding of MIG6 to an activating kinase domain interface. Nature 450(7170): 741–744. 10.1038/nature05998

[19] Jin N, Cho SN, Raso MG, et al. (2009) Mig-6 is required for appropriate lung development and to ensure normal adult lung homeostasis. Development (Cambridge, England) 136(19): 3347–3356. 10.1242/dev.032979

[20] Hopkins S, Linderoth E, Hantschel O, et al. (2012) Mig6 is a sensor of EGF receptor inactivation that directly activates c-Abl to induce apoptosis during epithelial homeostasis. Developmental cell 23(3): 547–559. 10.1016/j.devcel.2012.08.001

[21] Ferby I, Reschke M, Kudlacek O, et al. (2006) Mig6 is a negative regulator of EGF receptor-mediated skin morphogenesis and tumor formation. Nature medicine 12(5): 568–573. 10.1038/nm1401

[22] Staal B, Williams BO, Beier F, Vande Woude GF, Zhang YW (2014) Cartilage-specific deletion of Mig-6 results in osteoarthritis-like disorder with excessive articular chondrocyte proliferation. Proceedings of the National Academy of Sciences of the United States of America 111(7): 2590–2595. 10.1073/pnas.1400744111

[23] Pujar M, Vastrad B, Kavatagimath S, Vastrad C, Kotturshetti S (2022) Identification of candidate biomarkers and pathways associated with type 1 diabetes mellitus using bioinformatics analysis. Scientific Reports 12(1): 9157. 10.1038/s41598-022-13291-1

[24] Chen YC, Colvin ES, Griffin KE, Maier BF, Fueger PT (2014) Mig6 haploinsufficiency protects mice against streptozotocin-induced diabetes. Diabetologia 57(10): 2066–2075. 10.1007/s00125-014-3311-z

25. Council NR (2011) Guide for the Care and Use of Laboratory Animals: Eighth Edition. The National Academies Press, Washington, DC

[26] Zhang YW, Su Y, Lanning N, et al. (2005) Targeted disruption of Mig-6 in the mouse genome leads to early onset degenerative joint disease. Proceedings of the National Academy of Sciences of the United States of America 102(33): 11740–11745. 10.1073/pnas.0505171102

[27] Thorens B, Tarussio D, Maestro MA, Rovira M, Heikkila E, Ferrer J (2015) Ins1(Cre) knock-in mice for beta cell-specific gene recombination. Diabetologia 58(3): 558–565. 10.1007/s00125-014-3468-5

[28] Hingorani SR, Petricoin EF, Maitra A, et al. (2003) Preinvasive and invasive ductal pancreatic cancer and its early detection in the mouse. Cancer cell 4(6): 437–450

[29] Stull ND, Breite A, McCarthy R, Tersey SA, Mirmira RG (2012) Mouse islet of Langerhans isolation using a combination of purified collagenase and neutral protease. J Vis Exp(67). 10.3791/4137

[30] Chen YC, Colvin ES, Maier BF, Mirmira RG, Fueger PT (2013) Mitogen-inducible gene 6 triggers apoptosis and exacerbates ER stress-induced beta-cell death. Molecular endocrinology (Baltimore, Md) 27(1): 162–171. 10.1210/me.2012-1174

[31] Chen Y-C, Colvin ES, Fueger PT (2013) Mitogen-inducible gene 6 potentiates glucolipotoxicity-induced pancreatic beta cell death. The FASEB Journal 27(1_supplement): lb743-lb743. 10.1096/fasebj.27.1_supplement.lb743

[32] Colvin ES, Ma H-Y, Chen Y-C, Hernandez AM, Fueger PT (2013) Glucocorticoid-Induced Suppression of β-Cell Proliferation Is Mediated by Mig6. Endocrinology 154(3): 1039–1046. 10.1210/en.2012-1923

[33] Furman BL (2021) Streptozotocin-Induced Diabetic Models in Mice and Rats. Current Protocols 1(4): e78. https://doi.org/10.1002/cpz1.78

[34] Hahn M, van Krieken PP, Nord C, et al. (2020) Topologically selective islet vulnerability and self-sustained downregulation of markers for β-cell maturity in streptozotocin-induced diabetes. Communications Biology 3(1): 541. 10.1038/s42003-020-01243-2

[35] Lenzen S (2008) The mechanisms of alloxan-and streptozotocin-induced diabetes. Diabetologia 51(2): 216–226. 10.1007/s00125-007-0886-7

[36] Kim-Muller JY, Fan J, Kim YJR, et al. (2016) Aldehyde dehydrogenase 1a3 defines a subset of failing pancreatic β cells in diabetic mice. Nature Communications 7(1): 12631. 10.1038/ncomms12631

[37] Cinti F, Bouchi R, Kim-Muller JY, et al. (2016) Evidence of beta-cell dedifferentiation in human type 2 diabetes. Journal of Clinical Endocrinology and Metabolism 101: 1044–1054. 10.1210/jc.2015-2860

[38] Thorel F, Népote V, Avril I, et al. (2010) Conversion of adult pancreatic α-cells to β-cells after extreme β-cell loss. Nature 464: 1149. 10.1038/nature08894 https://www.nature.com/articles/nature08894#supplementary-information

[39] Gribben C, Lambert C, Messal HA, et al. (2021) Ductal Ngn3-expressing progenitors contribute to adult β cell neogenesis in the pancreas. Cell Stem Cell 28(11): 2000–2008.e2004. https://doi.org/10.1016/j.stem.2021.08.003

[40] Zhao H, Huang X, Liu Z, et al. (2021) Pre-existing beta cells but not progenitors contribute to new beta cells in the adult pancreas. Nature Metabolism 3(3): 352–365. 10.1038/s42255-021-00364-0

[41] Kopp JL, Dubois CL, Schaffer AE, et al. (2011) Sox9<sup>+</sup> ductal cells are multipotent progenitors throughout development but do not produce new endocrine cells in the normal or injured adult pancreas. Development (Cambridge, England) 138(4): 653

[42] Dor Y, Brown J, Martinez OI, Melton DA (2004) Adult pancreatic β-cells are formed by self-duplication rather than stem-cell differentiation. Nature 429(6987): 41–46. 10.1038/nature02520

[43] Ackermann AM, Gannon M (2007) Molecular regulation of pancreatic beta-cell mass development, maintenance, and expansion. Journal of molecular endocrinology 38(1-2): 193–206. 10.1677/jme-06-0053

[44] Teta M, Long SY, Wartschow LM, Rankin MM, Kushner JA (2005) Very Slow Turnover of β-Cells in Aged Adult Mice. Diabetes 54(9): 2557. 10.2337/diabetes.54.9.2557

[45] Wilcox CL, Terry NA, Walp ER, Lee RA, May CL (2013) Pancreatic α-Cell Specific Deletion of Mouse Arx Leads to α-Cell Identity Loss. PLOS ONE 8(6): e66214. 10.1371/journal.pone.0066214

[46] Sarnobat D, Charlotte Moffett R, Flatt PR, Irwin N, Tarasov AI (2022) GABA and insulin but not nicotinamide augment α-to β-cell transdifferentiation in insulin-deficient diabetic mice. Biochemical Pharmacology 199: 115019. https://doi.org/10.1016/j.bcp.2022.115019

[47] Maachi H, Fergusson G, Ethier M, et al. (2020) HB-EGF Signaling Is Required for Glucose-Induced Pancreatic β-Cell Proliferation in Rats. Diabetes 69(3): 369–380. 10.2337/db19-0643

[48] Suarez-Pinzon WL, Lakey JR, Brand SJ, Rabinovitch A (2005) Combination therapy with epidermal growth factor and gastrin induces neogenesis of human islet {beta}-cells from pancreatic duct cells and an increase in functional {beta}-cell mass. The Journal of clinical endocrinology and metabolism 90(6): 3401–3409. 10.1210/jc.2004-0761

[49] Tang S, Zhang M, Zeng S, et al. (2020) Reversal of autoimmunity by mixed chimerism enables reactivation of β cells and transdifferentiation of α cells in diabetic NOD mice. 117(49): 31219–31230. doi:10.1073/pnas.2012389117

[50] Chung C-H, Hao E, Piran R, Keinan E, Levine F (2010) Pancreatic β-Cell Neogenesis by Direct Conversion from Mature α-Cells. Stem Cells 28(9): 1630–1638. 10.1002/stem.482

[51] Thorel F, Népote V, Avril I, et al. (2010) Conversion of adult pancreatic α-cells to β-cells after extreme β-cell loss. Nature 464(7292): 1149–1154. 10.1038/nature08894

[52] Piran R, Lee SH, Li CR, Charbono A, Bradley LM, Levine F (2014) Pharmacological induction of pancreatic islet cell transdifferentiation: relevance to type I diabetes. Cell death & disease 5(7): e1357. 10.1038/cddis.2014.311

[53] Wang MY, Dean ED, Quittner-Strom E, et al. (2021) Glucagon blockade restores functional β-cell mass in type 1 diabetic mice and enhances function of human islets. Proceedings of the National Academy of Sciences of the United States of America 118(9). 10.1073/pnas.2022142118

[54] Tang S, Zhang M, Zeng S, et al. (2020) Reversal of autoimmunity by mixed chimerism enables reactivation of β cells and transdifferentiation of α cells in diabetic NOD mice. Proceedings of the National Academy of Sciences of the United States of America 117(49): 31219–31230. 10.1073/pnas.2012389117

[55] Xu D, Li C (2021) Gene 33/Mig6/ERRFI1, an Adapter Protein with Complex Functions in Cell Biology and Human Diseases. Cells 10(7). 10.3390/cells10071574

[56] Basavarajappa HD, Irimia JM, Bauer BM, Fueger PT (2023) The Adaptor Protein NumbL Is Involved in the Control of Glucolipotoxicity-Induced Pancreatic Beta Cell Apoptosis. 24(4): 3308

[57] Liu L, Xing L, Chen R, et al. (2021) Mitogen-Inducible Gene 6 Inhibits Angiogenesis by Binding to SHC1 and Suppressing Its Phosphorylation. Frontiers in cell and developmental biology 9: 634242. 10.3389/fcell.2021.634242

[58] Jiang X, Niu M, Chen D, et al. (2016) Inhibition of Cdc42 is essential for Mig-6 suppression of cell migration induced by EGF. Oncotarget 7(31): 49180–49193. 10.18632/oncotarget.10205

[59] Sasaki M, Terabayashi T, Weiss SM, Ferby I (2018) The Tumor Suppressor MIG6 Controls Mitotic Progression and the G2/M DNA Damage Checkpoint by Stabilizing the WEE1 Kinase. Cell reports 24(5): 1278–1289. 10.1016/j.celrep.2018.06.064

