## Supplementary Figures for "Pancreatic loss of Mig6 alters murine endocrine cell fate and protects functional beta cell mass in an STZ-induced model of diabetes"

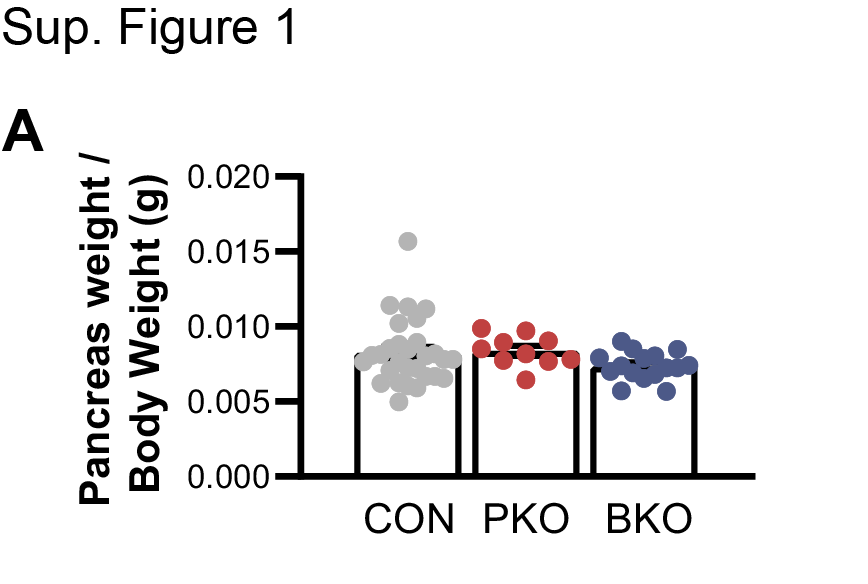

**Supplementary Figure 1: Mouse pancreas to body weight ratio**

Quantification of mouse pancreas weight divided by total body weight (g). n = 10-29 2-way ANOVA; p values: * <0.05 ** <0.01 *** <0.001; data reported as means ± SEM.

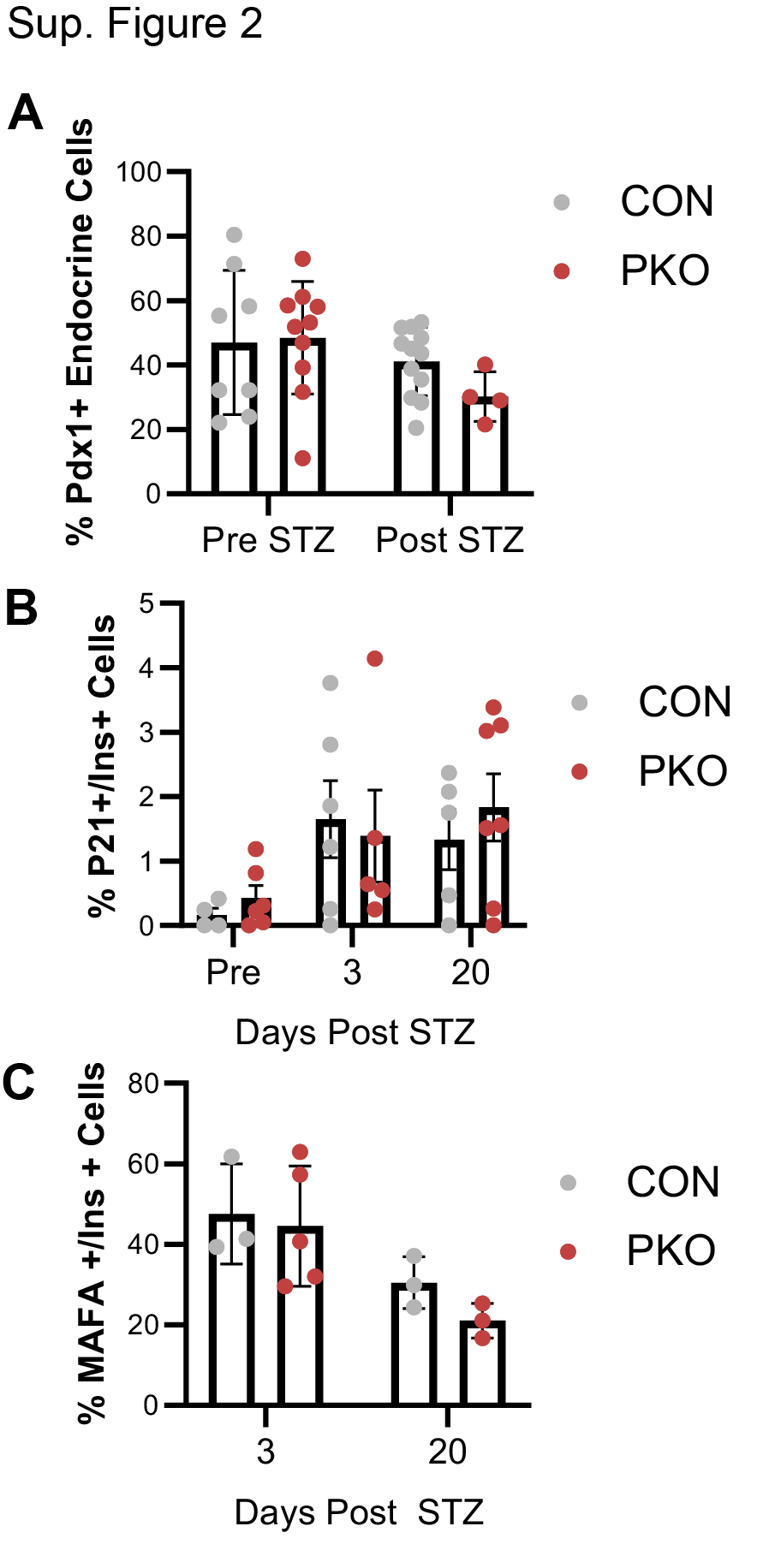

**Supplementary Figure 2: Additional markers of beta cell maturity and senescence following treatment with STZ**

**A.** Quantification of the percent of Pdx1 positive cells within islets as determined by QuPath both before and twenty days after STZ **B.** Quantification of p21 positive insulin positive senescent cells before STZ and then at three or twenty days following STZ. **C.** Quantification of MafA positive insulin positive mature beta cells at three and twenty days following treatment with STZ. n = 3-12, 2-way ANOVA or 2-tailed t-test; p values: * <0.05 ** <0.01 *** <0.001; data reported as means ± SEM.
